# A robust method for performing ^1^H-NMR lipidomics in embryonic zebrafish

**DOI:** 10.1101/2021.05.26.445770

**Authors:** Rhiannon S. Morgan, Gemma Walmsley, Mandy J. Peffers, Richard Barrett-Jolley, Marie M. Phelan

**Affiliations:** Institute of Life Course and Medical Sciences, University of Liverpool, Liverpool, UK; Institute of Systems, Molecular and Integrative Biology, University of Liverpool, Liverpool, UK

**Keywords:** lipidomics, ^1^H-NMR spectroscopy, zebrafish embryo

## Abstract

The embryonic zebrafish is an ideal system for lipid analyses with relevance to many areas of bioscience research, including biomarkers and therapeutics. Research in this area has been hampered by difficulties in extracting, identifying and quantifying lipids. We employed ^1^H-NMR at 700MHz to profile lipids in developing zebrafish embryos. The optimal method for lipidomics in embryonic zebrafish incorporated rapid lipid extraction using chloroform and an environment without oxygen depletion. Pools of 10 embryos gave the most acceptable signal-to-noise ratio, and the inclusion of chorions in the sample had no significant effect on lipid abundances. Embryos, bisected into cranial (head and yolk sac) and caudal (tail) regions, were compared by principal component analysis and analysis of variance. The lipid spectra (including lipid annotation) are available in the public repository MetaboLights (MTBLS2396).

## Introduction

Lipids are a diverse and abundant class of biomolecules within all living organisms [1]. Simple fats and oils provide energy and have important roles in processes such as signalling and membrane trafficking, whereas more complex, polar lipids such as sterols and phospholipids provide much needed structures to biological membranes [2]. The lipid bilayer is a key component of skeletal muscle membranes and here the lipids provide both structure and flexibility [3].

Disrupted lipid metabolism is a primary pathogenetic mechanism in a number of mammalian diseases including myopathies such as lipid storage myopathy and HACD1-associated centronuclear myopathy [4–6]. Despite the importance of lipids, the study of lipidomics is less advanced than that of genomics or proteomics. Gene and protein structures are linear alignments of code (base pairs, amino acids) whereas the structure of lipids is far more complex, making even the classification of these structure more difficult [1].

There are a vast number of techniques available to study lipid profiles, all with their own benefits and limitations. ^1^H-NMR is an important tool for lipidomic analyses, it is time efficient, highly reproducible and has been shown to identify a number of lipid classes in samples, including mouse liver [7]. NMR produces crowded spectra due to the lack of separation and so if separation is required mass spectrometry (MS) as a class of technique may be more appropriate [8]. MS is highly sensitive, however, the level of ionisation of lipids is variable and so lacks the reproducibility compared to NMR [8]. MS is destructive to samples and therefore complementary analyses cannot be performed on the same material, for these reasons’ NMR can be a desirable procedure for initial explorations [8].

Embryonic zebrafish are widely used to study development and genetic diseases, including several models for congenital myopathies [9–13]. Unlike mammals, zebrafish embryos develop externally and are therefore an isolated system for analysis of lipid metabolism during myogenesis [14]. Research in lipidomics (including biomarkers and therapeutics) has been hampered by difficulties in extracting, identifying and quantifying lipids. Previous studies have focused on the use of mass spectrometry techniques and while ^1^H-NMR has been conducted on zebrafish embryos this has mainly been to study polar/hydrophilic metabolites [15–19].

Robust methods for extraction and collection of ^1^H-NMR spectra are essential. Therefore, in this work, we established a working protocol that generates high-quality, reproducible spectra and confirmed our ability to detect lipidomic differences in developing embryos. In this study we aimed to carry out ^1^H-NMR lipid profiling in developing zebrafish embryos to establish a reproducible methodology for future studies, focusing on the protruding mouth stage (72 hours post fertilisation (hpf)).

## Materials and Methods

### Animals

Adult AB strain wild-type (WT) zebrafish were housed in a multi-rack aquarium system at the University of Liverpool. Zebrafish were maintained at 28.5 ± 0.5°C on a 12-hour light, dark cycle. Husbandry and collection of embryos involving zebrafish embryos were performed in accordance with both local guidelines (AWERB ref no. AWC0061) and within the UK Home Office Animals Scientific Procedures Act (1986), in a Home Office approved facility (University of Liverpool Establishment License X70548BEB). Adults were bred and embryos incubated in the dark at 28°C, in aquarium water and staged according to Kimmel et al., 1995 [20]. All experiments performed were exempt from ethical approval due to embryonic age as stipulated in ASPA guidelines.

### Embryonic Sample Collection

AB wildtype zebrafish (Danio rerio) embryos were collected into microcentrifuge tubes, and water was removed by careful pipetting before snap freezing in liquid nitrogen and storing at −80°C for a period of no more than 1 month. For all experiments except the chorion and development experiments, embryos aged 3 days post fertilisation (dpf) were used. For chorion experiments, 1dpf embryos were used and for dechorionated embryos the chorions were removed manually using a pair of fine watchmaker forceps (Dumont Laboratories) and discarded before snap freezing and storing as above. To bisect 3dpf embryos, embryos were washed in cold fish water (approximate temperature: 4°C) to shock and temporarily stop movement before using disposable scalpels (Swann-Morton) to bisect the embryos.

Whole embryos (all dpfs) were stored in pools of 10 embryos except where different embryo numbers required for the sample size experiments. Bisected embryos were stored in groups of 20 cranial or caudal portions per tube to counteract sample amount being decreased (Figure 1).

**Figure 1:**
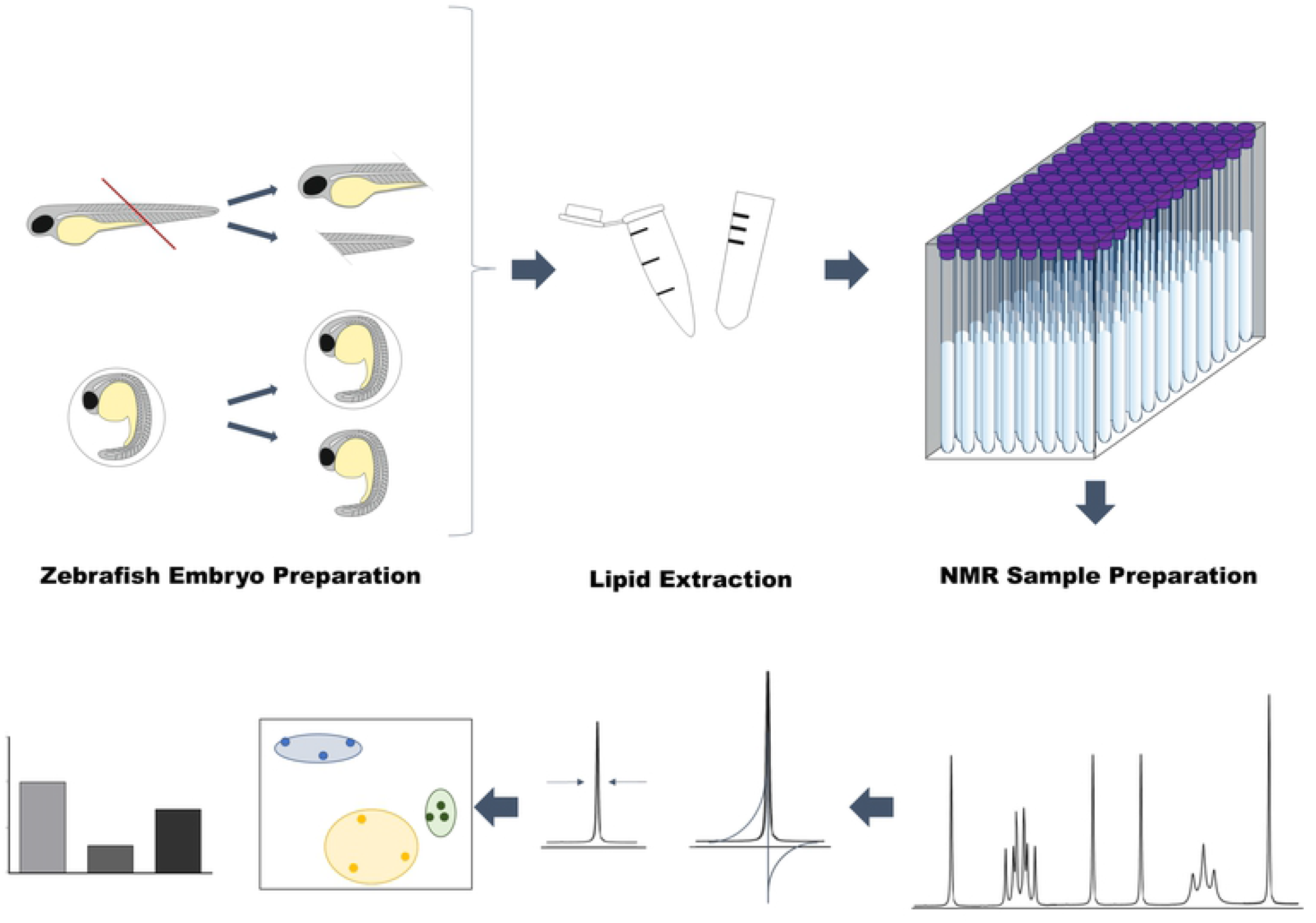
Schema of extraction protocol including zebrafish embryo preparation. Examples shown include the bisection of embryos 3 days post fertilisation (dpf) and removal of chorion from 1dpf embryos. After sample preparation lipids were extracted in either plastic Eppendorf tubes or glass mass spectrometry vials. Samples were run on a Bruker Avance III HD spectrometer with a TCI cryoprobe and chilled SampleJet autosampler with a field strength of 700.17MHz. Spectra were then processed and put through quality control measures (examples shown include phasing corrections and checking the chloroform peak width). Spectra were then binned appropriately using TameNMR and statistical analysis conducted using R.

### Lipid extraction

500μl of ice-cold chloroform (C_1_HC_l3_, Sigma Aldrich) was added to samples in microfuge tubes. Samples were then sonicated at 50KHz using a microprobe in three x 30 second intervals including a 30 second pause in between intervals. Samples were vortexed for one minute before incubating for 10 minutes at 4°C prior to centrifugation for 10 minutes at 21,500g and 4°C. The supernatant was transferred to a fresh microcentrifuge tube or glass via and snap frozen in liquid nitrogen before lyophilisation. Chloroform is a highly volatile chemical and is known to react with plastic, for this reason lipids were also extracted in glass mass spectrometry vials [21]. After plastic was deemed to be most suitable, lipids were extracted in both normoxic and hypoxic conditions under low oxygen levels (constant nitrogen stream) to determine if the oxidation of lipids would produce an observable effect on spectra [22] and to evaluate the optimum extraction method for ^1^H-NMR analysis.

### Sample preparation

Lyophilised samples were resuspended in 200μl of 99.8% deuterated chloroform (C2HCl3, Sigma Aldrich). Samples were vortexed for 10 seconds and centrifugated at 21500g, 4°C for one minute. The supernatant was then transferred using a glass Pasteur pipette to 3mm (outer diameter) NMR tubes.

### NMR Spectra acquisition

Samples were acquired on a Bruker Avance III HD spectrometer with a TCI cryoprobe and chilled SampleJet autosampler with a field strength of 700.17MHz. Spectra acquisition and processing was carried out using Topspin 3.1 software. Spectra were acquired and pre-processed according to standard metabolomics practices by an NMR spectroscopist at the Shared Research Facility for NMR metabolomics [14]. Briefly 1D ^1^H-NMR NOESY standard vendor pulse sequence (noesgppr1d - all parameters constant between samples) was used to acquire with a 25ppm spectral width and 256 scans (thirty-minute experiment) at 15°C to offset the volatility of the chloroform. Pre-processing of spectra proceeded using automated standard vendor routines (apk0.noe - Fourier transform, phasing and window function) and spectra were aligned using the residual C_1_HC_l3_ peak at 7.26ppm.

### Spectra inclusion criteria

Strict quality control (QC) was conducted on all spectra. Phasing of baselines was checked and manually corrected when required. The reference peak (C_1_HC_l3_) line width was measured at half height to ensure it was a single peak of <1.1Hz. Any samples that failed QC were acquired again on the spectrometer up to three times.

### Spectra processing

Using a representative set of spectra, a pattern file defining bin boundaries was generated for all peak positions. Where possible peaks were identified using and in-house Avanti Library or annotated according to published identities [23,24]. The pattern file was validated using the TameNMR Galaxy toolkit (https://github.com/PGB-LIV/tameNMR) and then used to integrate the spectral peaks data into numerous ‘bins’ for statistical analysis. Spectra were also subsetted for head-group analysis by omission of bins (1-112) attributed to hydrocarbon chain identities from published ^1^H-NMR studies [7, 24].

### Statistical analysis

Data was normalised by Probabilistic Quotient Normalisation (PQN) and auto scaled (each bin was mean centred followed division by the standard deviation for that bin) before performing multivariate (PCA) and univariate one-way ANOVA analyses. ANOVA was followed by Post-hoc analysis using Tukey’s Honest Significant Difference method for multiple comparisons. Cluster separations for PCA were scored and then we evaluated the probability that the separation seen was due to chance. This was performed using a permutation method within r package ‘ClusterSignificance’ [25]. Statistical analysis was conducted using in-house scripts implemented using the mixOmics package in R (r-project.org) [26].

## Results

Sample collection was based on Beckonert et al., (2007) and Folch (1957) with the extraction procedure simplified to focus solely on lipids and reduce sample losses (omitted two-phase extraction) (for entire workflow see Figure 1) [27, 28].

### Optimisation of sample size

Sample size experiments determined 10 embryos was sufficient to produce ^1^H-NMR spectra with 181 bins (Figure 2, Table 1). Figure 2 shows sections of the spectra corresponding to the majority of polar headgroups and the lipophilic hydrocarbon chain, with spectra from 1x embryo, 2x, 5x and 10x embryos at 3 days post fertilisation. Annotated bins for phosphatidylcholine (PC), triacylglyceride (TG) and total-cholesterol (TC) were selected due to their relevance in zebrafish development [9]. These bins show increasing signal in the spectra with increased embryo numbers within the sample. We calculated signal-to-noise ratios to establish whether bins exceeded both the limits of detection (LOD) and quantification (LOQ) (Table 1). Table 1 shows that certain bins did not always meet the LOQ or LOD (bins 118, 137 and 151) so these would not be appropriate bins to use for lipid quantification therefore other bins representing the same lipid classes should be selected, for example bin 150 was selected to quantify triacylglycerides over that of bin 151 (Table 1).

**Figure 2:**
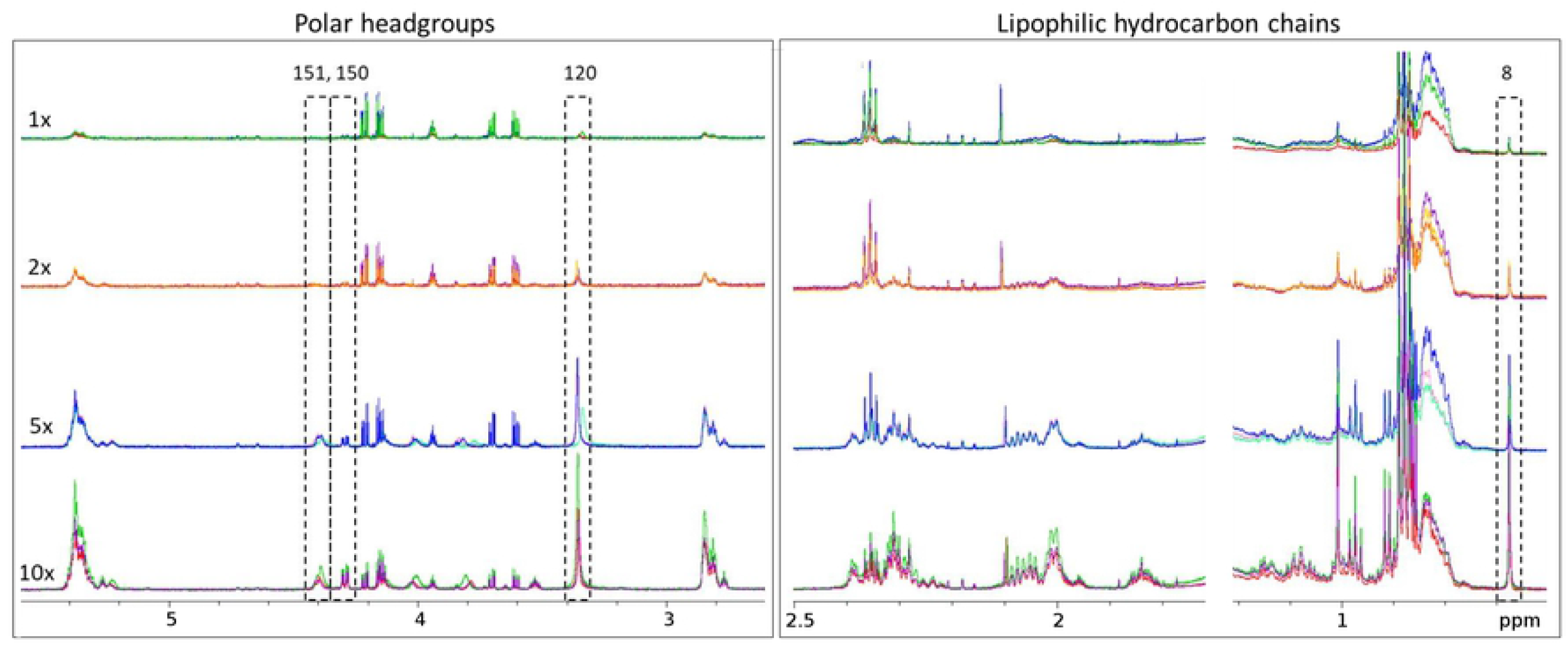
1D ^1^H-NMR spectra showing the polar headgroups and hydrocarbon chain portions of ZF extracts. 1D ^1^H-NMR spectra (Topspin) showing the polar headgroups and hydrocarbon chain portions of the spectra in sample sizes of 1, 2, 5 and 10 embryos at 3 days post fertilisation (n=3).

**Table 1:** A sample of the annotated spectral peaks and their mean signal-to-noise ratios for specific lipid bins corresponding to specific lipid classes.

|  |  |  |  |  |  |  |  |  |  |  |
| --- | --- | --- | --- | --- | --- | --- | --- | --- | --- | --- |
| Bin number | **8** | 20 | 118 | **120** | 123 | 136 | 137 | **150** | 151 | 152 |
| Bin ID | **Total Cholesterol** | Free Cholesterol | Lysophosphatidylcholine | **Phosphatidylcholine** | Free Cholesterol | Phospholipids | Phospholipids | **Triacylglycerides** | Triacylglycerides | Phospholipids |
| Chorion 1dpf (10)* | **2029.36** | 10271.17 | 28.25 | **276.16** | 182.91 | 84.41 | 210.12 | **1422.89** | 103.21 | 82.03 |
| no-chorion 1dpf (10)* | **2188.83** | 11043.17 | 39.61 | **301.31** | 190.96 | 600.87 | 72.16 | **1900.87** | 98.51 | 132.49 |
| 1dpf (10)* | **1586.26** | 8876.64 | 10.73 | **380.22** | 148.83 | 121.15 | 211.43 | **1241.07** | 129.81 | 93.87 |
| 2dpf (10)* | **477.39** | 2598.88 | 4.25 | **103.80** | 42.08 | 31.17 | 30.02 | **335.91** | 34.53 | 29.88 |
| 3dpf (1)* | **158.07** | 2478.67 | 0.31 | **25.22** | 18.61 | 80.56 | 10.77 | **138.01** | 8.09 | 169.51 |
| 3dpf (2)* | **159.16** | 1719.28 | -0.36 | **26.93** | 13.71 | 25.66 | 9.46 | **109.19** | 7.13 | 75.57 |
| 3dpf (5)* | **328.62** | 2173.93 | 2.09 | **71.78** | 27.18 | 30.79 | 15.82 | **200.41** | 24.09 | 63.24 |
| 3dpf (10)* | **785.24** | 4419.27 | 5.55 | **107.54** | 68.53 | 53.32 | 66.23 | **489.48** | 36.57 | 56.49 |
| Cranial (20) | **2626.67** | 13571.41 | 22.97 | **532.87** | 225.33 | 168.95 | 407.86 | **1824.24** | 190.55 | 121.12 |
| Caudal (20) | **235.23** | 1768.09 | -0.26 | **6.63** | 16.86 | 23.38 | 1.3 | **130.14** | 3.22 | 49.09 |
| Eppendorf with N2 (10) | **412.12** | 2088.55 | 0.84 | **43.39** | 34.16 | 173.29 | 5.03 | **287.21** | 15.46 | 13.81 |
| Glass with N2(10) | **709.42** | 4138.22 | 10.84 | **116.75** | 58.65 | 298.71 | 36.41 | **508.96** | 41.31 | 23.54 |

Bins that failed the LOD (limit of detection) threshold of 3:1 are highlighted in yellow and those failing the LOQ (limit of quantification) threshold of 10:1 in
blue. Spectral bins 8, 120 and 150 (in bold) were selected for further analysis whereas bins 118, 137 and 151 are examples of bins that did not meet the LOD
certain sample types. Dpf=days post fertilisation.
\*whole embryo extraction in Eppendorf without O<sub>2</sub> depletion

### The effect of embryonic dechorionation

Removal of the chorion revealed no benefit with regards to reduced variation and samples did not cluster separately (*p*-value=0.894) (Figure 3A). ANOVA results identified no significant changes in lipid abundance in any of the bins (adjusted *p*-value <0.05) between the two samples (Figure 3A).

**Figure 3:**
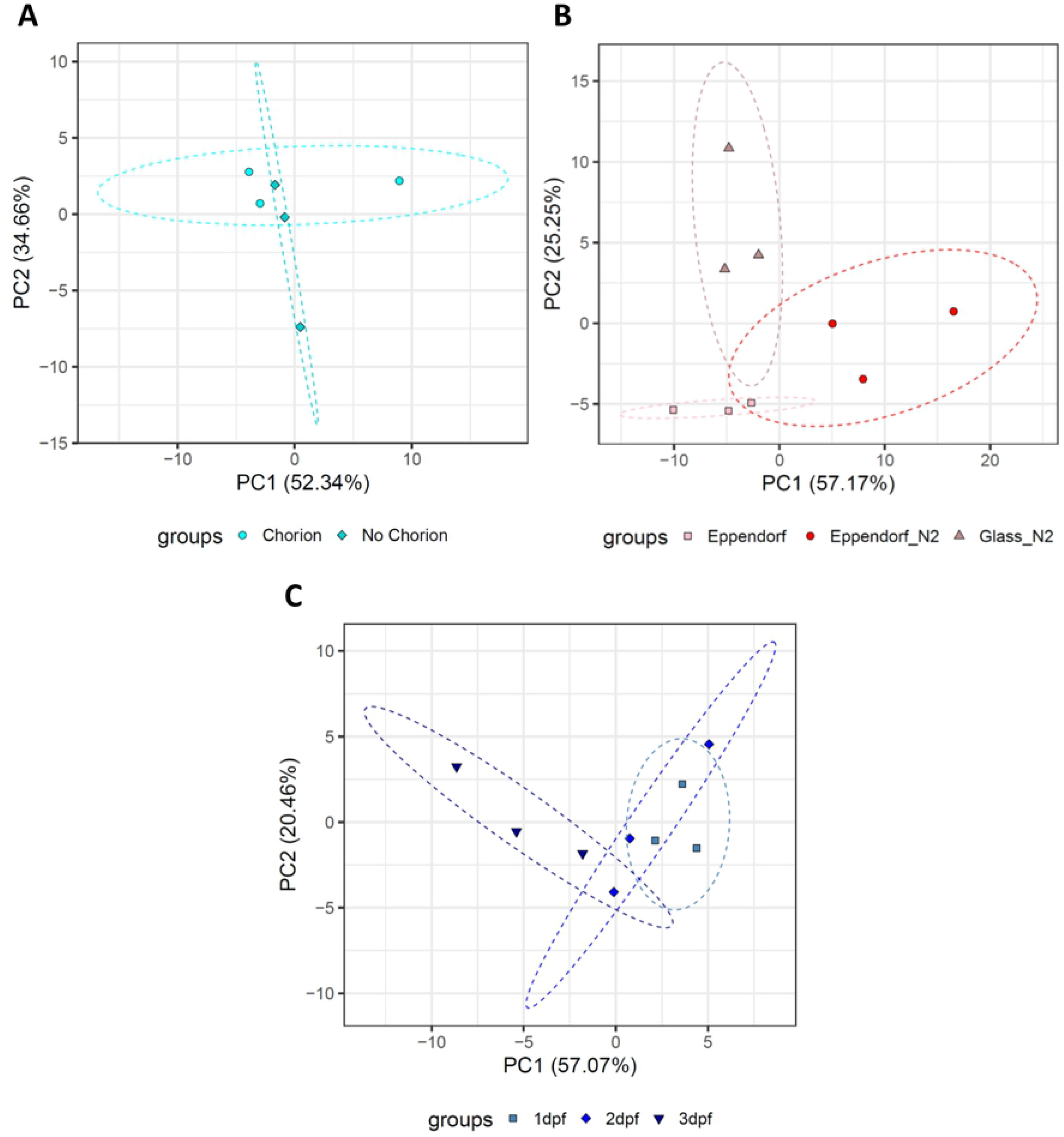
Effects of sample processing, extraction and developmental stage. (A) principal component analysis (PCA) scores of 1 day post fertilisation (dpf) embryos with and without chorions (n=3). (B) PCA scores for 3dpf embryonic lipids extracted in either glass vials with nitrogen gas (N_2_), or Eppendorf tubes with or without N2 (n=3). (C) PCA scores of embryos during development (n=3).

### The effect of extraction environment

Spectra were compared from extractions performed in glass vials and Eppendorfs with or without the depletion of oxygen by use of nitrogen gas (N2). The addition of sample preparation in a N2 environment exhibited a larger variance within the group spectra for both glass and Eppendorf extracts in low O2 environments with respect to extracts performed in Eppendorfs extracted without O_2_ depletion (Figure 3B). ANOVA results found 67 significant bins between the three extraction environments (Figure 3B).

### Lipidomic evaluation of developmental stages and bisected embryos

The lipidome of embryos during development was also compared using unsupervised multivariate principal component analysis (PCA), with 3dpf embryos clustering almost separately from 1dpf (*p*-value=0.09) embryos but with 2dpf overlapping 1 and 3dpf embryos (*p*-values=0.116 and 0.906 respectively) (Figure 3C). ANOVA results identified 10 significant bins (adjusted *p*-value <0.05) between the developmental stages, with all 10 of these differing between 1 and 3dpf (Figure 3C).

The marginal separation of clusters seen between developmental stages (Figure 3C) led us to assess how much influence the yolk sac may be having in the spectra. Spectra acquired on dissected cranial (head and yolk sacs) and caudal (tails) sections gave clear separation with cranial sections and whole embryos clustered together (*p*-value=0.349), but separately from the caudal sections (caudal versus whole *p*-value=0.026, caudal versus cranial *p*-value=0.001) which are predominantly muscle (Figure 4A). The PCA loadings for whole embryo (Figure 4B) show the variance between the cranial sections and whole versus the caudal section. Univariate analysis confirmed that these differences could be attributed to a large number of bins (132 bins of 181 adjusted *p*-value <0.05), of which three bins of biological interest have been annotated in red (Figure 4B). Analysis using the section of the spectra attributed to the head group regions of lipids showed similar clustering patterns with caudal sections separated out and also separation between whole embryos and cranial sections (Figure 4C). The PCA loadings indicated that the variance between cranial and caudal sections was dominated by peaks attributed to this lipid headgroup region (bins 113-181) (Figure 4D).

**Figure 4:**
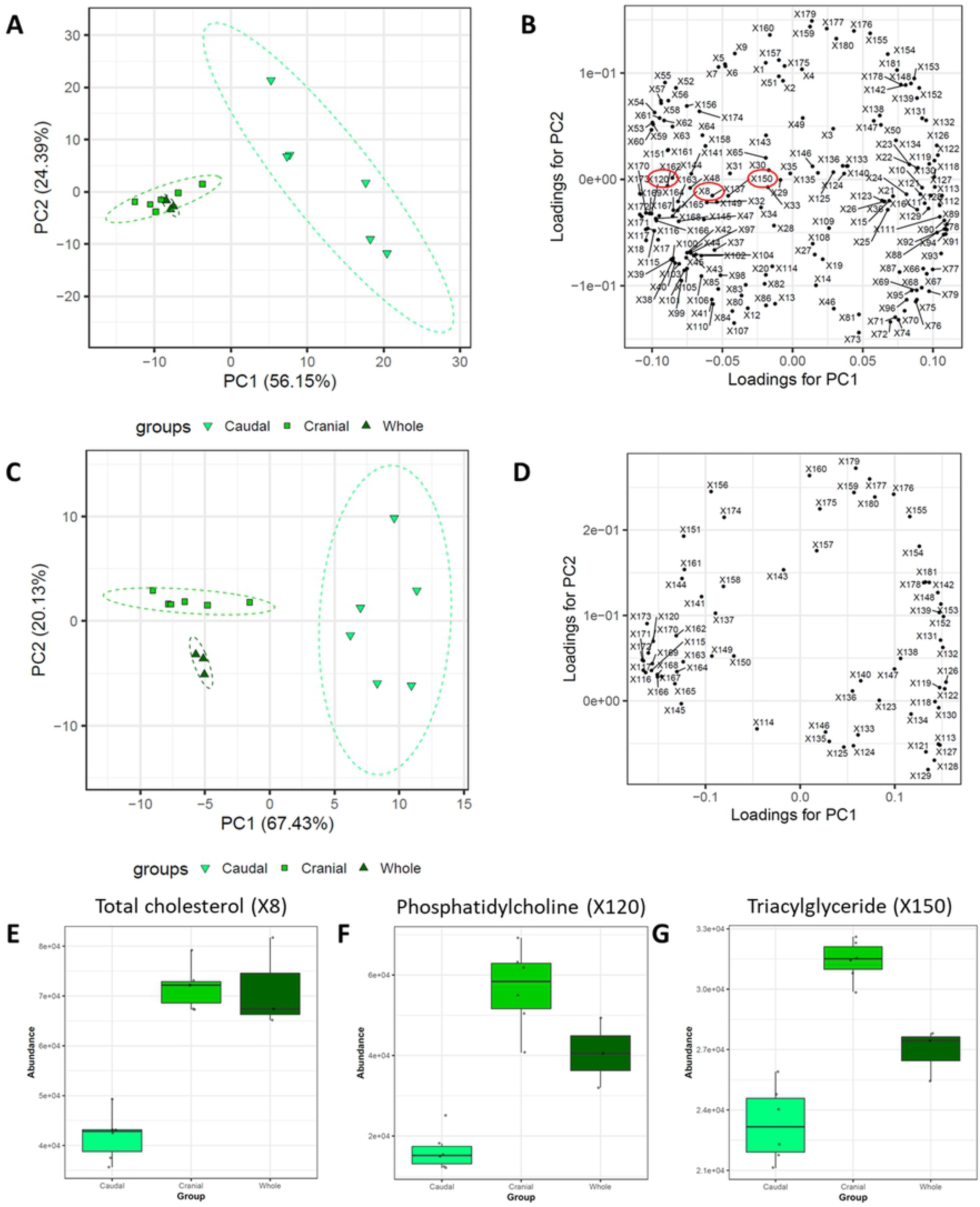
The effect of bisecting embryos prior to lipid analysis. (A) Principal component analysis (PCA) scores for whole embryos versus bisected cranial/caudal sections and (B) corresponding PCA loadings (n=3 whole, n=6 cranial/caudal sections). (C) PCA scores for whole embryos versus bisected cranial/caudal sections using only the main headgroup region (bins 113-181) of the spectra to further identify peaks of interest and remove variation created by sample loss) and (D) corresponding PCA loadings (n=3 whole, n=6 cranial/caudal sections). (E-G) Representative boxplots for total cholesterol (TC) (E), phosphatidylcholine (PC) (F) and triacylgylcerides (TG), these bins are annotated on Figure 3B (G). Adjusted *p*-values for E-F all equal < 0.05, ANOVA. Tukey’s multiple comparison showed Cranial-Caudal and Whole-Caudal to be significant, adjusted *p*-values < 0.05 (E) and all three comparisons to be significant in (F) and (G) (adjusted *p*-values < 0.05). Dotted lines (A&C) indicate the 95% confidence interval.

The annotated bins from 3B are in Figures 3E-G showing the relative abundance of representative peaks for total cholesterol (TC), phosphatidylcholine (PC) and triacylglycerides (TG) respectively. ANOVAs for these 3 bins found them to be significantly changed as a whole (*p*<0.05) with relative abundance of these bins lower in the caudal sections. Tukey’s multiple comparisons found TC to be significantly lower in caudal sections than in either the cranial sections or whole embryos (*p*<0.05) and all three groups caudal, cranial and whole significantly different for PC and TG (*p*<0.05).

## Discussion

Lipidomics in embryonic zebrafish is an emerging field and has potential for numerous applications with far-reaching impact. This work provides a robust methodology for ^1^H-NMR lipidomics using embryonic zebrafish from which reproducible, informative spectra can be produced.

An increase in embryo number increased signal-to-noise ratio and allowed for the acquisition of a quantitative lipid spectra (Figure 2). Pools of 10 embryos provided a spectra in which all bins tested passed both the limit of detection and limit of quantification (Figure 2). Previous studies have used large numbers of zebrafish embryos (~100 embryos) for ^1^H-NMR [18,19]. Whilst zebrafish can lay clutches of several hundred embryos when housed in optimal conditions [29], pools of 100 are typically not experimentally feasible due to time constraints from commonly used techniques such as microinjection. Therefore, our finding that it is possible to use pools of 10 embryos for ^1^H-NMR lipidomic analysis is important for studies of genetically modified embryos.

Dechorionation was deemed to be an unnecessary step because there was no significant difference in the lipidome between samples with and without chorions (Figure 3A). This was an expected result as chorions are proteinaceous structures and not lipid based [30]. We were also able to conclude that retaining the chorions benefits spectral consistency – a probable reason for the increased variability is the possibility of an induced metabolic change in the sample. Embryos hatch from their chorions between 2 – 3 days post fertilisation (dpf) therefore when analysing different developmental changes, we recommend collection of embryos without including this labour-intensive step [20].

We found the most robust method to extract lipids was rapid extraction in plastic Eppendorf tubes without the depletion of oxygen by use of nitrogen gas (N_2_). The use of increased variation seen in the PCA plots (Figure 3B) which led to a number of significant changes within the ^1^H-NMR spectra (Figure 3D). We propose that the additional air-flow induced through N_2_ rich working may have been a factor in increased variance in the lipid metabolome and outweighs the benefit of oxygen depletion on the lipidome in this study.

There was less cluster separation between embryos of different developmental stages than expected (Figure 3C) and we hypothesised this may be due to the yolk sac masking any effects seen from within the rest of the embryo due to its high lipid composition [15]. To test this, we bisected the embryos into cranial (head and yolk sac) and caudal (tail) regions and found the caudal sections segregated completely, thus confirming our hypothesis (Figure 4A & C). Fraher et al., 2016 determined that at 0 hours post fertilisation (hpf) when the embryo consists of solely yolk sac, the most abundant classes of lipid were: cholesterol (TC) followed by phosphatidylcholines (PC) and triacylglycerides (TG). In our data these three lipid classes are significantly lower in the caudal samples and highest the cranial region (in which the yolk sac is the greatest proportion of sample) and then slightly lower in whole embryos. Therefore, bisection or removal of the yolk sac may be necessary in some situations including detailed analysis of the lipidome in a particular tissue (tail muscle for example) or evaluation of low abundance lipid classes.

In conclusion we have developed a robust method for analysis of lipid profiles of zebrafish embryos by 1D ^1^H-NMR. Furthermore, we anticipate this protocol could broaden the scope of ^1^H NMR lipidomic analysis by providing a template for establishing lipid analysis methods in other model organisms by ^1^H-NMR spectroscopy.

## Additional Information

### Abbreviations

NMR: nuclear magnetic resonance
dpf: days post fertilisation
hpf: hours postfertilisation
WT: wild type
QC: quality control
ANOVA: analysis of variance
PCA: principal component analysis
PQN: probabilistic quotient normalisation
PC: phosphatidylcholine
TG: triacylglyceride
TC: total-cholesterol
FC: free-cholesterol
LOD: limit-of-detection
LOQ: limit-of-quantification

## Institutional Review Board Statement

Ethical review and approval were waived for this study, due to embryonic age exemption as stipulated in ASPA guidelines.

## Data availability

The spectra and annotation are available at the MetaboLights public repository MetaboLights ID: MTBLS2396 [31].

## Author Contributions

Rhiannon Morgan: Conceptualization, Methodology, Validation, Formal analysis, Investigation, Writing - Original Draft, Writing - Review & Editing, Visualization, Funding acquisition. Gemma Walmsley: Conceptualization, Methodology, Validation, Resources, Writing - Review & Editing, Supervision, Funding acquisition. Mandy Peffers: Methodology, Writing - Review & Editing, Supervision. Richard Barrett-Jolley: Methodology, Writing - Review & Editing, Supervision. Marie Phelan: Conceptualization, Methodology, Validation, Formal analysis, Resources, Data Curation, Writing - Original Draft, Writing - Review & Editing, Visualization, Supervision, Project administration.

All authors have read and agreed to the published version of the manuscript.

## References

1. Fahy E, Cotter D, Sud M, Subramaniam S. Lipid classification, structures and tools. Biochim Biophys Acta 2011. 1811(11):637–647. DOI: 10.1016/j.bbalip.2011.06.009.

2. Lehninger AL, Nelson DL, Cox MM. Lehninger principles of biochemistry. 2000. 3rd edition. Worth Publishers, New York.

3. van Meer G, Voelker DR, Feigenson GW. Membrane lipids: where they are and how they behave. Nat Rev Mol Cell Bio. 2008. 9(2):112–124. DOI: 10.1038/nrm2330.

4. Bruno C, Dimauro S. Lipid storage myopathies. Curr Opin Neurol. 2008. 21(5):601–606. DOI: 10.1097/WCO.0b013e32830dd5a6.

5. Blondelle J, Ohno Y, Gache V, Guyot S, Storck S, Blanchard-Gutton N, et al. HACD1, a regulator of membrane composition and fluidity, promotes myoblast fusion and skeletal muscle growth. J Mol Cell Bio. 2015. 7(5):429–40. DOI: 10.1093/jmcb/mjv049.

6. Walmsley GL, Blot S, Venner K, Sewry C, Laporte J, Blondelle J. et al. Progressive Structural Defects in Canine Centronuclear Myopathy Indicate a Role for HACD1 in Maintaining Skeletal Muscle Membrane Systems. Am J Pathol. 2015. 187(2): 441–456. 10.1016/j.ajpath.2016.10.002.

7. Amiel A, Tremblay-Franco M, Gautier R, Ducheix S, Montagner A, Polizzi A. Proton NMR Enables the Absolute Quantification of Aqueous Metabolites and Lipid Classes in Unique Mouse Liver Samples. Metabolites. 2019. 10(1):9. DOI: 10.3390/metabo10010009.

8. Li J, Vosegaard T, Guo Z. Applications of nuclear magnetic resonance in lipid analyses: An emerging powerful tool for lipidomics studies. Progress in Lipid Research. 2017. 68: 37–56. DOI: 10.1016/j.plipres.2017.09.003.

9. Dowling JJ, Vreede AP, Low SE, Gibbs EM, Kuwada JY, Bonnemann CG, et al. Loss of myotubularin function results in T-tubule disorganization in zebrafish and human myotubular myopathy. PLoS Genetics. 2009. 5(2):e1000372. DOI: 10.1371/journal.pgen.1000372.

10. Dowling JJ, Low SE, Busta AS, Feldman EL. Zebrafish MTMR14 is required for excitation–contraction coupling, developmental motor function and the regulation of autophagy. Human Molecular Genetics. 2010. 19(13):2668–2681. DOI: 10.1093/hmg/ddq153.

11. Gibbs EM, Davidson AE, Trickey-Glassman A, Backus C, Hong Y, Sakowski SA, et al. Two Dynamin-2 Genes Are Required for Normal Zebrafish Development. PloS One. 2013. 8(2), e55888. DOI: 10.1007/s13311-018-00686-0.

12. Sabha N, Volpatti JR, Gonorazky H, Reifler A, Davidson AE, Li X, et al. PIK3C2B inhibition improves function and prolongs survival in myotubular myopathy animal models. Journal of Clinical Investigation. 2016. 126(9):3613–3625. DOI: 10.1172/JCI86841.

13. Smith LL, Gupta VA, Beggs AH. Bridging integrator 1 (Bin1) deficiency in zebrafish results in centronuclear myopathy. Human Molecular Genetics. 2014. 23(13):3566–3578. DOI: 10.1093/hmg/ddu067.

14. Guyon JR, Steffen LS, Howell MH, Pusack TJ, Lawrence C, Kunkel LM. Modeling human muscle disease in zebrafish. Biochimica et Biophysica Acta. 2007. 1772(2):205–15. DOI: 10.1016/j.bbadis.2006.07.003.

15. Fraher D, Sanigorski A, Mellett NA, Meikle PJ, Sinclair AJ, Gibert Y. Zebrafish Embryonic Lipidomic Analysis Reveals that the Yolk Cell Is Metabolically Active in Processing Lipid. Cell Reports. 2016. 14(6):1317–1329. DOI: 10.1016/j.celrep.2016.01.016.

16. Xu M, Legradi J, Leonards P. Evaluation of LC-MS and LC×LC-MS in analysis of zebrafish embryo samples for comprehensive lipid profiling. Anal Bioanal Chem. 2020. 412:4313–4325. DOI: 10.1038/s41598-017-17409-8.

17. Zhang W, Song Y, Chai T, Liao G, Zhang L, Jia Q, et al. J. Lipidomics perturbations in the brain of adult zebrafish (Danio rerio) after exposure to chiral ibuprofen. Sci Total Environ. 2020. 713:136565. DOI: 10.1016/j.scitotenv.2020.136565.

18. Roy U, Conklin L, Schiller J. Metabolic profiling of zebrafish (Danio rerio) embryos by NMR spectroscopy reveals multifaceted toxicity of β-methylamino-L-alanine (BMAA). Sci Rep. 2017. 7:17305. DOI: 10.1038/s41598-017-17409-8.

19. Zuberi Z, Eeza MNH, Matysik J, Berry JP, Alia A. NMR-Based Metabolic Profiles of Intact Zebrafish Embryos Exposed to Aflatoxin B1 Recapitulates Hepatotoxicity and Supports Possible Neurotoxicity. Toxins. 2019. 11(5):258. DOI: 10.3390/toxins11050258.

20. Kimmel CB, Ballard WW, Kimmel SR, Ullmann B, Schilling TF. Stages of embryonic development of the zebrafish. Developmental Dynamics. 1995. 203(3):253–310. DOI: 10.1002/aja.1002030302.

21. Norman S Radin. Lipids and Related Protocols. Humana Press, 1988. US. DOI: 10.1385/0-89603-124-1:1.

22. Crowe T, Crowe T, Johnson L, White P. Impact of extraction method on yield of oxidation products from oxidized and unoxidized walnuts. Journal of Oil & Fat Industries. 2002. 79. 453–456. DOI: 10.1007/s11746-002-0505-7.

23. Sparling. M. L. Analysis of mixed lipid extracts using 1H NMR spectra. Bioinformatics. 1990. 6(1):29–42.

24. Tynkkynen T. 1H NMR Analysis of Serum Lipids. PhD Dissertation, 2002. University of Eastern Finland, Finland, ISBN: 978-952-61-0838-4.

25. Serviss JT, Gådin JR, Eriksson P, Folkersen L, Grandér D. ClusterSignificance: a bioconductor package facilitating statistical analysis of class cluster separations in dimensionality reduced data. Bioinformatics. 2017. 33(9):3126–3128. DOI: 10.1093/bioinformatics/btx393.

26. Lê Cao KA, Costello ME, Lakis VA, Bartolo F, Chua XY. MixMC: A Multivariate Statistical Framework to Gain Insight into Microbial Communities. PLOS ONE. 2016. 11(8):e0160169. DOI:10.1371/journal.pone.0160169.

27. Beckonert O, Keun HC, Ebbels TM, Bundy J, Holmes E, Lindon JC, et al. Metabolic profiling, metabolomic and metabonomic procedures for NMR spectroscopy of urine, plasma, serum and tissue extracts. Nature Protocols. 2007. 2(11):2692–703. DOI: 10.1038/nprot.2007.376.

28. Folch J, Lees M, Sloane Stanely. GH. A simple method for the isolation and purification of total lipides from animal tissues. J Biol Chem. 1957. May, 226(1):497–509. PMID: 13428781.

29. Nasiadka A, Clark MD. Zebrafish breeding in the laboratory environment. ILAR Journal. 2012. 53(2):161–168. DOI: 10.1093/ilar.53.2.161.

30. Bonsignorio D, Perego L, Del Giacco L, Cotelli F. Structure and macromolecular composition of the zebrafish egg chorion. Zygote. 1996. 4(2):101–108. DOI: 10.1017/s0967199400002975.

31. Haug K, Salek RM, Conesa P, Hastings J, de Matos P, Rijnbeek M, Mahendraker T. et al. MetaboLights - an open-access general-purpose repository for metabolomics studies and associated meta-data. Nucleic Acids Res. 2013. 41(Database issue):D781–6. DOI: 10.1093/nar/gks1004.

